# Inhibition of proprotein convertase subtilisin-like kexin type 9 (PCSK9) potentiates anti-angiogenic therapy in colorectal cancer liver metastases

**DOI:** 10.1101/2023.05.29.542731

**Authors:** Miran Rada, Andrew R. Reynolds, Anthoula Lazaris, Nabil Seidah, Peter Metrakos

## Abstract

Colorectal cancer liver metastatic (CRCLM) tumours present as two main histopathological growth patterns (HGPs) including desmoplastic HGP (DHGP) and replacement HGP (RHGP). The DHGP tumours obtain their blood supply by sprouting angiogenesis, whereas the RHGP tumours utilize an alternative vascularisation known as vessel co-option. In vessel co-option, the cancer cells hijack the mature sinusoidal vessels to obtain blood supply. Vessel co-option has been reported as an acquired mechanism of resistance to anti-angiogenic treatment in CRCLM. Herein, we showed that inhibiting proprotein convertase subtilisin-like kexin type 9 (PCSK9) via clinically approved PCSK9-neutralizing antibody (Evolocumab) can boost the response of vessel co-opting tumours to anti-angiogenic therapy. Mechanistically, we found that PCSK9 inhibition downregulates runt related transcription factor-1 (RUNX1) expression levels in CRCLM cancer cells in vivo, which its expression positively correlates with the development of vessel co-option. Collectively, these results suggest that inhibiting PCSK9 is a promising way to improve the efficacy of anti-angiogenic therapy against vessel co-opting tumours in CRCLM.

## Introduction

Colorectal cancer (CRC) is the third most common cancer and the second deadliest cancer (1), which accounts for approximately 10% of cancer-related death in men and women (2). The liver is the most common site of metastases of colorectal cancer, and approximately 50% of patients diagnosed with CRC develop liver metastases (LM) during the course of their disease(3). Surgical resection improves survival by up to 50% and could be a potentially curative treatment for CRCLM patients (4). However, only 15-20% of CRCLM patients are eligible for surgery resection(5). The rest of the patients are treated with chemotherapy and targeted therapies such as anti-angiogenic agents (e.g. Bevacizumab)(6). However, acquired resistance against these treatments frequently occurs that resulting in treatment failure and cancer recurrence (7).

Colorectal cancer liver metastases (CRCLMs) present with two common histopathological growth patterns (HGPs): Desmoplastic HGP (DHGP) and replacement HGP (RHGP) (8). The fundamental histopathological feature of DHGP lesions is the presence of a desmoplastic rim at the tumour periphery that separates cancer cells from liver parenchyma(7). However, the desmoplastic rim is absent in RHGP lesions, and the cancer cells infiltrate through adjacent liver plates (9,10). Moreover, angiogenesis is the predominant vascularization in DHGP tumours (7). However, vessel co-option is the prevalent vascularization in RHGP tumours, in which the cancer cells co-opt the pre-existing vessels to obtain the blood supply (11,12). Previous investigations reported vessel co-option as acquired resistance to anti-angiogenic therapy in various cancers including CRCLM, hepatocellular carcinoma (HCC) and glioblastoma (13).

We previously identified various molecules that regulate the generation of vessel co-opting tumours in CRCLM, such as runt related transcription factor-1 (RUNX1) (14) and actin related protein 2/3 complex (ARP2/3) (7). Accordingly, cancer cells in vessel co-opting CRCLM tumours express higher levels of RUNX1, which is a transcriptional regulator of ARP2/3 (14). Both RUNX1 and ARP2/3 facilitate the development of vessel co-opting tumours via increasing cancer cells motility, which allows the cancer cells to infiltrate through liver parenchyma around tumour nest (7).

Various studies indicated a positive association between the risk of developing cancer and the elevation of serum cholesterol levels (15). Cholesterol is an important lipid for supporting cellular homeostasis (16). Low-density lipoprotein (LDL) is the primary carrier of cholesterol in the blood to deliver cholesterol to peripheral and liver cells, and LDL receptor (LDLR) is the major determinant of LDL uptake by the liver (17). The levels of LDL in the blood are controlled by proprotein convertase subtilisin-like kexin type 9 (PCSK9), which targets LDLR for lysosomal degradation (18,19). Evolocumab is the monoclonal antibody against PCSK9 that binds to the circulating PCSK9 protein, inhibiting it from binding to the LDLR (20). Evolocumab has been approved for the treatment of hypercholesterolaemia (21). Recently, various preclinical studies suggested evolocumab treatment to increase the efficacy of anti-cancer agents, such as doxorubicin and trastuzumab (22). Moreover, Liu et al. (23) have reported that evolocumab potentiated immune checkpoint therapy against various types of tumours in vivo. However, the role of PCSK9 in increasing efficacy of anti-angiogenic agents against vessel co-option is largely unknown, specifically in CRCLM.

In this manuscript, we showed that treatment with anti-PCSK9 (evolocumab) significantly potentiates the function of anti-angiogenic therapy against vessel co-option tumours. Our data also suggested that treating tumours with evolocumab dramatically downregulate RUNX1 expression in the cancer cells in vivo.

## Materials and methods

### Cell culture

Mouse colorectal cancer (MC38) cells were cultured in DMEM (Wisent Inc., #319-005-CL) supplemented with 10% FBS (Wisent Inc., #085-150) and 1× penicillin/streptomycin (Wisent Inc., 450-201-EL). All cells were cultured at 37□°C with 5% CO2.

### Immunohistochemical Staining

We performed immunohistochemical staining for formalin-fixed paraffin-embedded (FFPE) specimens as described in previous publications (24,25). We performed staining using RUNX1 antibody 1:200 (LS Bio, #LS-C353932).

### Xenograft experiments

CRCLM tumours were generated in C57B/6 mice via intrasplenic injection of mouse colorectal cancer (MC38) cells (5*10^5^ cells/ mice) as described in previous publications (26). Five days after intrasplenic injection, the mice were divided into four groups (n=5 mice per group) after intrasplenic injection. The mice in the first group were injected with a vehicle alone (Control). The mice in the second group were treated with intraperitoneal injection of 2.5 mg per kg of VEGF-A inhibitory antibody (B20-4.1.1) every five days. The mice in the third group were injected intraperitoneally with 200μg of evolocumab (also called Repatha; Amgen Manufacturing Limited) every two days. The fourth group was treated with B20-4.1.1 combined with evolocumab.

### Statistical reproducibility

Statistical analysis was performed using GraphPad Prism software version 9.0 (GraphPad Software, La Jolla, CA, USA) software. For the overall survival data, the Log-Rank test was used to determine the statistical significance. P-values of <0.05 were considered to be significant.

## Results

### Treatment with evolocumab increases the efficacy of anti-angiogenic therapy against vessel co-opting tumours

We previously reported that treating tumours with evolocumab attenuates the generation of vessel co-opting tumours in CRCLM (25). However, the role of evolocumab in increasing the efficacy of anti-angiogenic (anti-VEGF) therapy in CRCLM is unclear. Thus, we decided to perform intrasplenic injection of mouse colorectal cancer (MC38) cells into C57B/6 mice. Of note, previous studies all CRCLM tumours that generated from MC38 cancer cells are vessel co-opting tumours (26). We divided the mice as follows: a. control; b. treated with anti-VEGF; c. treated with evolocumab; d. treated with anti-VEGF combined with evolocumab. The experiment was ended 50 days after intrasplenic injection. The mice were sacrificed once they became sick. The mice in the control group were sacrificed 17-19 days after intrasplenic injection (Figure 1a). The treated mice with anti-angiogenic therapy were sacrificed 18-20 days after intrasplenic injection. However, the treated mice with evolocumab were sacrificed 42-50 days after days after intrasplenic injection. Intriguingly, all mice that were treated with a combination of anti-angiogenic therapy and evolocumab were healthy even 50 days after intrasplenic injection.

**Figure 1.**
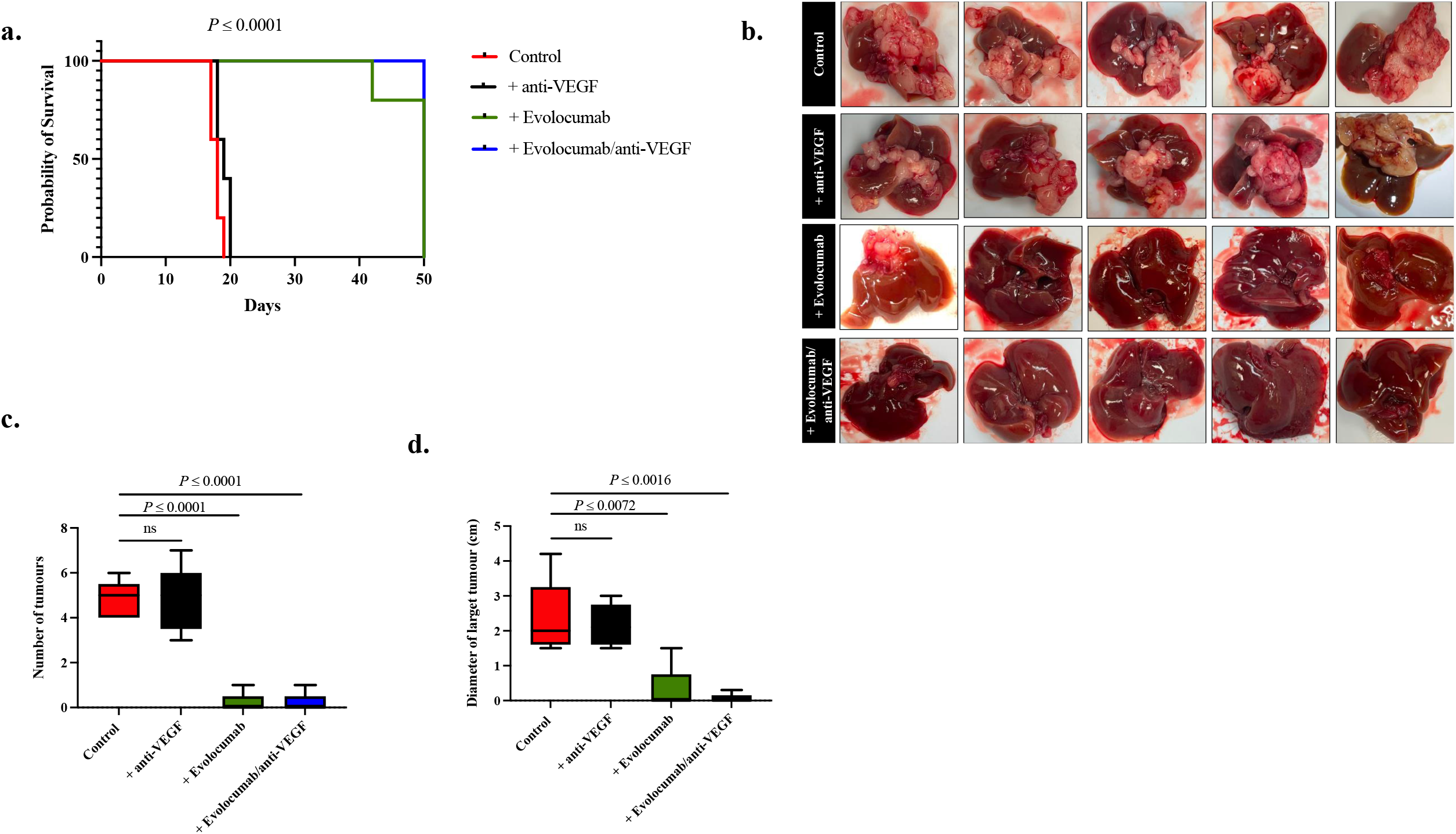
Treatment with evolocumab potentiates anti-angiogenic therapy in CRCLM tumours in vivo. **a**. Kaplan– Meier survival curves showing overall survival of mice with CRCLM tumiurs in the presence or absence of treatment with evolocumab and/or anti-angiogenic therapy. **b**. Images of the harvested liver of the mice that received intra-splenic injections of MC-38 cells. **c**. Represent the number of the generated liver metastatic tumours. **d**. Represent the size and diameter of the generated liver metastatic tumours.

After harvesting the livers, we found metastatic tumour in the liver of all mice from group a and b, while each of group c and d had only one mice with generated CRCLM (Figure 1b). Interestingly, the number and size of the generated tumours in group c and d were significantly lower compared to group a and b (Figure 1c and 1d). These results suggest that evolocumab reduces the number and size of the CRCLM in vivo, as well as sensitize CRCLM tumours to anti-angiogenic therapy.

### Treatment with evolocumab attenuates the expression levels of RUNX1 in cancer cells in vivo

We previously found that inhibition of RUNX1 in cancer cells attenuates their capability for proliferation and motility, as well as impairs the development of vessel co-opting RHGP tumours in CRCLM (14). To determine the correlation between evolocumab treatment and RUNX1 expression in CRCLM tumours. We performed immunohistochemical staining using anti-RUNX1 antibody for CRCLM sections that were generated from control (non-treated) and evolocumab-treated mice. Our data showed dramatic reduction of RUNX1 expression in the cancer cells that were treated with evolocumab compared to the cancer cells that were not treated with evolocumab (Figure 2). These data suggest inverse correlations between RUNX1 expression in CRCLM cancer cells and evolocumab treatment.

**Figure 2.**
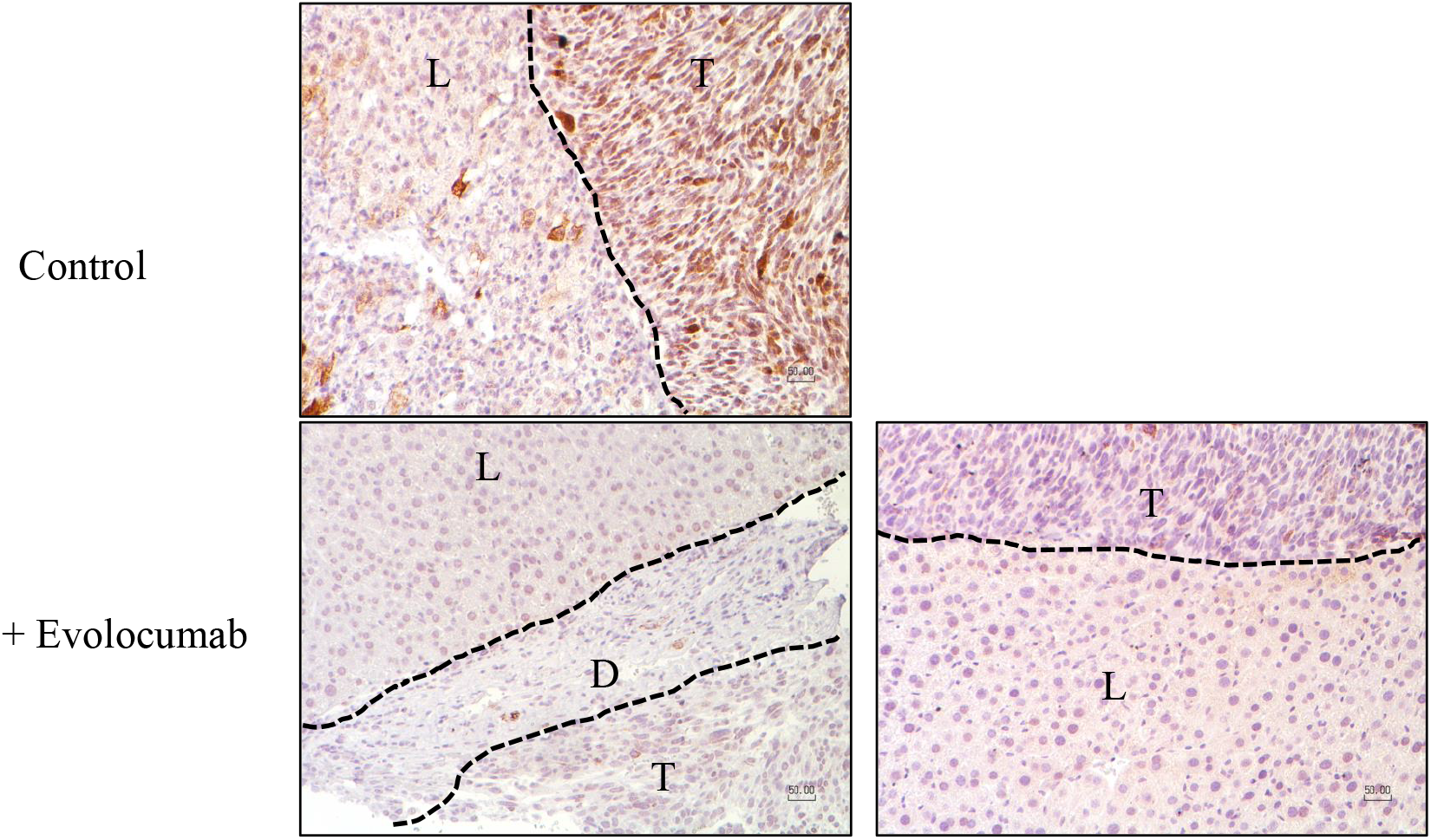
Treatment with evolocumab reduces RUNX1 expression in CRCLM cancer cells in vivo. Immunohistochemical staining using anti-RUNX1 antibody to determine the presence of RUNX1. The CRCLM sections were generated by intrasplenic injection of MC38 cancer cells into C57B/6 mice.

## Discussion

Vessel co-option is a non-angiogenic vascularization method that allows the tumours to further their growth and spread by hijacking the pre-existing vasculature (10). Therefore, the tumours that utilize vessel co-option vascularization are resistant to anti-angiogenic therapy (7). Vessel co-option is a predominant vascularization in CRCLM (7,9). We previously reported that CRCLM patients with predominantly angiogenic metastasis receiving neoadjuvant anti-angiogenic plus chemotherapy have significantly higher 5-year overall survival compared to the patients with co-opting tumours who have received the same treatment (7). These data suggest that the function of anti-angiogenic therapy against vessel co-opting CRCLM tumour is limited. Therefore, finding new agents that potentiate anti-angiogenic function in treating vessel co-option has become of utmost importance. Herein, we found evolocumab potentiates the impact of anti-angiogenic therapy in CRCLM.

The upregulation of PCSK9 expression has been linked to poor prognosis in various cancers, such as HCC (27), CRC (28), prostate cancer (29), and ovarian cancer (30). The involvement of PCSK9 in tumour resistance against anti-cancer agents has been investigated previously. Recent studies reported that anti-PCSK9 antibodies including evolocumab and alirocumab increase the effect of immune checkpoint therapy in vivo (23). However, the role of anti-PCSK9 antibodies in resistance to anti-angiogenic therapy is poorly understood. Our data suggested that inhibition of PCSK9 potentiates anti-angiogenic therapy in vivo. In agreement with our results, another study has linked PCSK9 overexpression to resistance against sorafenib, an anti-angiogenic agent, in HCC (31).

Our data suggested that treatment with evolocumab alone is also effective and attenuates the development of metastatic CRC tumour metastases in vivo. Wang et al. (32)have reported that knockdown of PCSK9 attenuates colon cancer cell proliferation, migration, and invasion and suppressed tumor metastasis. Of note, we previously reported that RUNX1 plays a positive regulator of cancer cell proliferation, migration, and invasion in CRCLM (14). In the current study, we noticed dramatic downregulation of RUNX1 expression in CRCLM cancer cells upon evolocumab treatment in vivo. These data suggest the molecular mechanisms by which evolcumab attenuates the generation of CRCLM. Indeed, further investigations are required to identify the mediator(s) between PCSK9 and RUNX1.

## Conclusions

Our preclinical data demonstrated that PCSK9 inhibition increases the efficacy of anti-angiogenic therapy CRCLM. Moreover, our results proposed the molecular mechanisms by which PCSK9 inhibition attenuates the development of CRCLM in vivo.

## Author Contributions

M.R., A.L., A.R., N.S. and P.M. co-conceived the study. M.R. performed cell culturing, immunohistochemistry, xenograft experiments, mice treatment, data curation, writing and preparation of the manuscript. P.M. funding acquisition.

## Institutional Review Board Statement

The study was approved by the McGill University Health Centre Institutional Review Board.

## Data Availability Statement

The data presented in this study are available within the article.

## Acknowledgments

The authors would like to acknowledge the support provided by Dana Massaro and Ken Verdoni Liver Metastases Research Fellowship. We also thank Genenthech for providing VEGF-A inhibitory antibody (B20-4.1.1).

